# Exploring Directed Connectivity Patterns of OPM-MEG with Multivariate Transfer Entropy

**DOI:** 10.1101/2024.02.22.581475

**Authors:** Mangor Pedersen, Geet Vashista, Dion Henare, Paul F. Sowman

## Abstract

We explored the directed connectivity of magnetoencephalography data recorded via Optically Pumped Magnetometers (OPM-MEG). Ten healthy adult participants were scanned twice while watching a movie in an OPM-MEG system. Beamformer source localisation was employed to obtain source time series within 62 cortical brain regions. Multivariate transfer entropy (mTE) was used to quantify directed connectivity. Our results suggest that out-degree connectivity (node-to-network) was stronger than in-degree (network-to-node). Out-degree display a more heavy-tailed distribution, indicative of brain hubs, whereas in-degree nodes had a distribution closer to a normal distribution. Although in-degree and out-degree nodes were positively correlated, we observed that out-degree connectivity was stronger in somatomotor and attention networks than in other resting-state networks. However, in- and out-degree were both inversely related to the physical distance between nodes, suggesting that directed connectivity is stronger between short-distanced nodes. As OPM-MEG research is likely to expand substantially in the future, we propose that directed connectivity measures may provide additional insights into the underlying mechanisms of brain communication.

## Background

Rapid advances in quantum physics are changing the field of neuroscience. A prime example is the recent advent of the OPM-MEG scanner (Boto et al., 2018). OPM-MEG records fast neural activity at the millisecond scale, overcoming common temporal limitations of techniques such as fMRI (Boto et al., 2019; Hill et al., 2020, 2022; Jas et al., 2021; Mellor et al., 2021; Roberts et al., 2019). Additionally, it is less susceptible to volume conduction, overcoming many of the source localisation and functional connectivity limitations of EEG (Brookes et al., 2022). OPM-MEG works by measuring magnetic fields generated by currents in assemblies of neurons in the brain, enabled by recording OPM-MEG in a shielded room that deflects any interference from the earth’s magnetic field (Holmes et al., 2022; Iivanainen et al., 2019; Seymour et al., 2022). In a room with virtually no magnetic field from the earth, OPMs use light/laser to cause a state transition in magnetically susceptible Alkali vapours, enabling detection of the magnetic field from ensembles of neurons when they are active in the brain (Tierney et al., 2019). While Alkali-based sensors are used in this study, it is worth noting that helium-4 OPM sensors are also emerging (Fourcault et al., 2021). An advantage of OPM-MEG is that its magnetic field recording ability is less affected by confounds and artefacts commonly associated with traditional neuroscience equipment (e.g., functional MRI), such as head motion, allowing users to move during a scan. Such advances open up research possibilities in challenging cohorts such as children and clinical populations, including people with seizures and movement disorders (Feys et al., 2022; Mellor et al., 2021; Pedersen et al., 2022; Vivekananda et al., 2020).

Pioneering OPM-MEG research has demonstrated that it can maintain comparable performance to SQUID-based MEG even while participants make naturalistic movements (Botos et al. 2018), outperforming other wearable imaging like EEG (Boto et al., 2019). Therefore, recent reports have leveraged OPM-MEG to derive functional connectivity patterns using correlation-based connectivity methods (Boto et al., 2021) and established good test-retest reliability (Rier et al., 2023). However, no work has been done to date exploring directed connectivity as a method with OPM-MEG, as it enables a bi-directional investigation of information transfer between nodes.

Some of the most common directed connectivity measures include the model-based Dynamic Causal Modelling Field (Friston et al., 2003), as well as the Granger Causality Field (Granger, 1969) and Transfer Entropy (Schreiber, 2000). We used Transfer Entropy in this study due to its advantageous multivariate and non-parametric properties (Novelli et al., 2019). The primary objective of our research study was to investigate various aspects of directed connectivity utilising OPM-MEG combined with Multivariate Transfer Entropy (mTE - Novelli et al., 2019; Wollstadt et al., 2019). This statistical measure based on information theory quantifies the information transferred from one time series to another, encompassing the input and output of mTE (as introduced by Schreiber, 2000). Unlike other measures, mTE connectivity matrices are asymmetric, allowing us to quantify the information going *to and from* different regions with a specific temporal distance – i.e., at one particular temporal lag. Consequently, mTE provides a framework for investigating directed connectivity between brain regions, shedding light on the specific communication pathways within the brain.

The main objective of this study was to investigate the performance of mTE-based directed connectivity measures using OPM-MEG data, extending previous work from non-directed connectivity findings in OPM-MEG (Boto et al., 2021; Rier et al., 2023) as well as other methodologies (Betzel et al., 2013; Bullmore & Sporns, 2009; Sporns, 2013; van den Heuvel et al., 2008). Our study had several specific aims to assess the performance of mTE. Given that the main advantage of directed connectivity (here estimated with mTE) is the ability to estimate incoming and outgoing nodal connectivity information, the main objective of this study was to explore the similarities and differences of in- and out-degree, in the context of OPM-MEG. We specifically aimed to test whether in/out-degree differs between pre-defined ‘resting-state’ networks.

Additionally, we aimed to examine the relationship between Transfer Entropy and the physical distance between brain sources. Our study provides insights into the capabilities and characteristics of mTE in estimating directed connectivity, by leveraging the potential of OPM-MEG technology.

## Methods

### Subjects

We used data from previous work (Rier et al., 2023), which is openly available in a Zenodo repository (https://zenodo.org/records/7477061) under a Creative Commons Attribution 4.0 International licence allowing re-use. The study was conducted and ethically approved at the University of Nottingham. No local ethics approval was needed for this study. In brief, this study had 10 participants, 6 males and 4 females, all right-handed, who provided written informed consent to participate in the experiment. The age range of the participants was 31 ± 8 years (mean ± standard deviation across subjects).

### OPM-MEG

In a magnetically shielded room, participants underwent two scanning sessions in an OPM-MEG system with 168 triaxial channels (Brookes et al., 2021). Before the OPM-MEG recording, a field-mapping and nulling procedure (Rea et al., 2021) was conducted to control the background magnetic field. Participants viewed a 600-second clip from the movie “Dog Day Afternoon” used in published papers (Haufe et al., 2018; Honey et al., 2012). Each subject underwent two scanning sessions with the same clip for both scans. The sensor helmet was not removed between scans to ensure a single co-registration of sensor geometry to brain anatomy, thereby minimising co-registration error. The time gap between the two scanning runs was approximately 1-2 minutes. This procedure ensured the accuracy and consistency of the magnetic field measurements throughout the experiment.

We pre-processed the data using MATLAB scripts (v2023a, Natick, Massachusetts) from https://github.com/LukasRier/Rier2022_OPM_connectome_test-retest/. Every participant also underwent an MRI scan, which was performed using a Phillips Ingenia 3T MRI scanner with an MPRAGE sequence and a 1-mm isotropic resolution. An anatomical MRI and a 3D in-line surface representation (Einscan H, SHINING 3D, Hangzhou, China) were used to co-register accurate sensor locations. A single-shell forward model was computed using a head model built using the MNI-152 template MRI with a 1 mm isotropic voxel resolution (Nolte, 2003). The source location was presented in a pseudo-Z picture reconstructed using a Linearly Constrained Minimum Variance Beamforming beamformer. We calculated the Beamformed signal within 62 cortical nodes (based on the AAL parcellation mask – Tzourio-Mazoyer et al., 2002). We only used cortical nodes due to the lower signal-to-noise ratio of deep sources with MEG (Boto et al., 2019). We applied a notch filter at 50Hz before additionally band-pass filtering the OPM-MEG time series between 1 and 48 Hz data using a 4th-order Butterworth Filter. More information on the OPM-MEG system used in this study can be found in Rier et al. (2023).

### Multivariate Transfer Entropy (mTE)

mTE aims to model the system by identifying a minimal set (***S*)** of sources that collectively contribute to the target’s next state (Novelli et al., 2019; Novelli & Lizier, 2021). After finding a subset of random variables as a candidates set, we estimate the conditional entropy between the node target and identified sources in the past set of lag targets. For mTE, the goal for each target is to identify the smallest number of sources past set (***X***^***S***^_**<*t***_,), which includes a non-uniform embedding of a target’s set previous state (γ_***t***_), that maximizes the collective entropy to target (**γ**^***S***^ _**<*t***_).

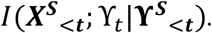

Here, *I* is conditional mutual information and multivariate variables are in **bold**.

In this approach from Novelli et al., (2019), a greedy approach is used to iteratively select candidate variables from a set of lagged variables from the past of each target and source. The greedy algorithm has several steps: (1) Evaluating the transfer entropy of all possible source nodes to a target note (2) Select for inclusion a source that accounts for the largest amount of incremental information about the next time step of the target node (3) repeat steps 1 and 2 until no additions can be made or all possible sources are included. Computational complexity is further addressed using set lags to restrict the field of possible time steps in the past likely contributing sources. The minimum lag in this study is 1 time point, and the maximum lag is 10 time points. A minimum statistic test was used to remove redundant sources in selecting past variables as part of the greedy algorithm. Final source selection was conducted with an Omnibus test that evaluates each calculated TE source against a null distribution with 200 randomisations in the embedding space under the assumption that no information is being transferred between signals. mTE was computed with a Gaussian Estimator using the IDTxl toolbox (https://github.com/pwollstadt/IDTxl).

We observed that over 80% of all Transfer Entropy connections had short lags within 1 or 2-time points with a heavy-tailed distribution towards a lag of 10-time points (see supplementary materials 1). Since mTE employs an in-built statistical procedure pruning connections in the network (Novelli et al., 2019), we observed that there were slight differences in network density between subjects (network density session 1 = 9.5% ± 2.2; and network density session 2 = 9.8% ± 2.3). Only binary networks were used in this work, which means that the lagged information is not taken into account in the network-specific analyses.

### Graph theoretic metrics

We calculated graph measures with directed information by computing the In-Degree (incoming information to nodes) and Out-Degree (outgoing information to nodes). See Rubinov & Sporns (2010), for more information. In-Degree is calculated as 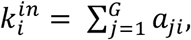 and Out-Degree is calculated as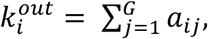. With G being the size of the network (here 62 nodes or sources), as the sum of lags are measured in the network (*a*_*ij*_ and *a*_*ji*_, respectively). Note that *a*_*ij*_ and *a*_*ji*_ are not equal and result in directional connectivity information (see Figure 1).

**Figure 1:**
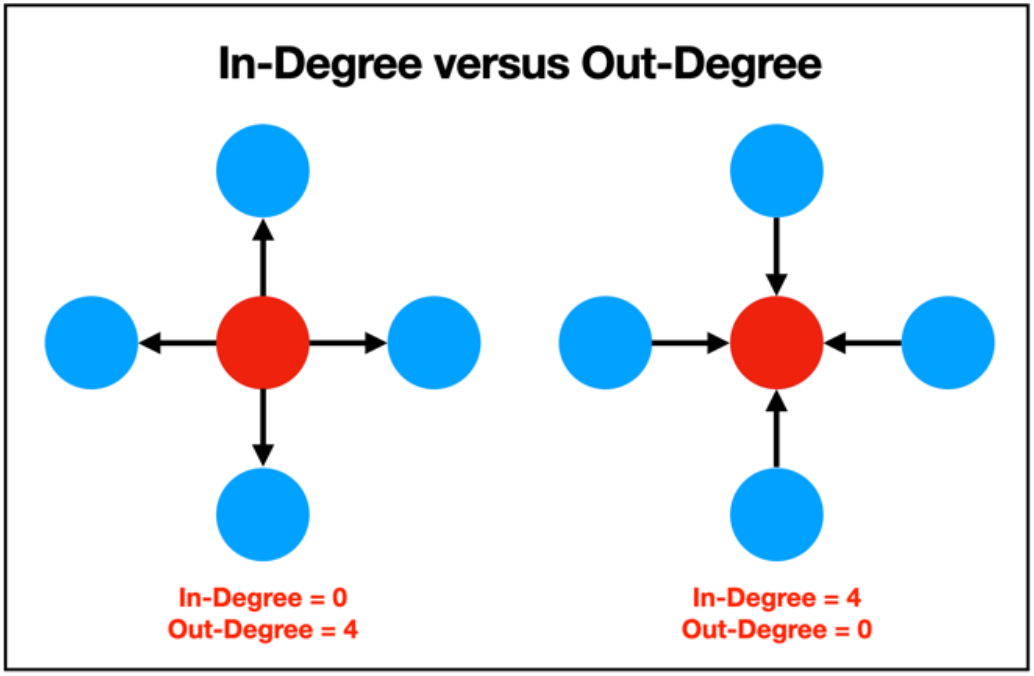
Two toy networks describing the difference between In- and Out-Degree. Left network: a network with low In-Degree and high Out-Degree. Right: a network with high In-Degree and low Out-Degree.

### Statistical analysis

To estimate in- and out-degree differences between resting-state networks, we employed a 1×6 ANOVA, and multiple comparisons testing was assessed with Tukey’s test. Spearman’s rho correlations were used to quantify the relationships between In- and Out-Degree, as well as the physical length between nodes.

## Results

### Relationship between Transfer Entropy derived In- and Out-Degree

Firstly, we observed that Out-Degree nodes display a more heavy-tailed distribution, indicative of brain hubs, whereas in-degree nodes had a distribution closer to normal (Figure 2A). In- and Out-Degree nodes were positively correlated despite their distribution differences. Session #1: Spearman’s ρ(61) = 0.59, *p* < 0.001. Session #2: Spearman’s ρ(61) = 0.42, *p* < 0.001 (see Figure 2B). These findings suggest that nodes with higher In-Degree also tend to have higher Out-Degree, reflecting interconnectedness within the network. Despite the significant correlation between In- and Out-Degree, we did observe brain-specific differences between In- and Out-Degree nodes. To this end, we calculated the mean difference between In- and Out-Degree nodes and found that the frontal cortex has higher In-Degree and nodes across somatosensory and posterior cortices have higher Out-Degree (Figure 2C).

**Figure 2:**
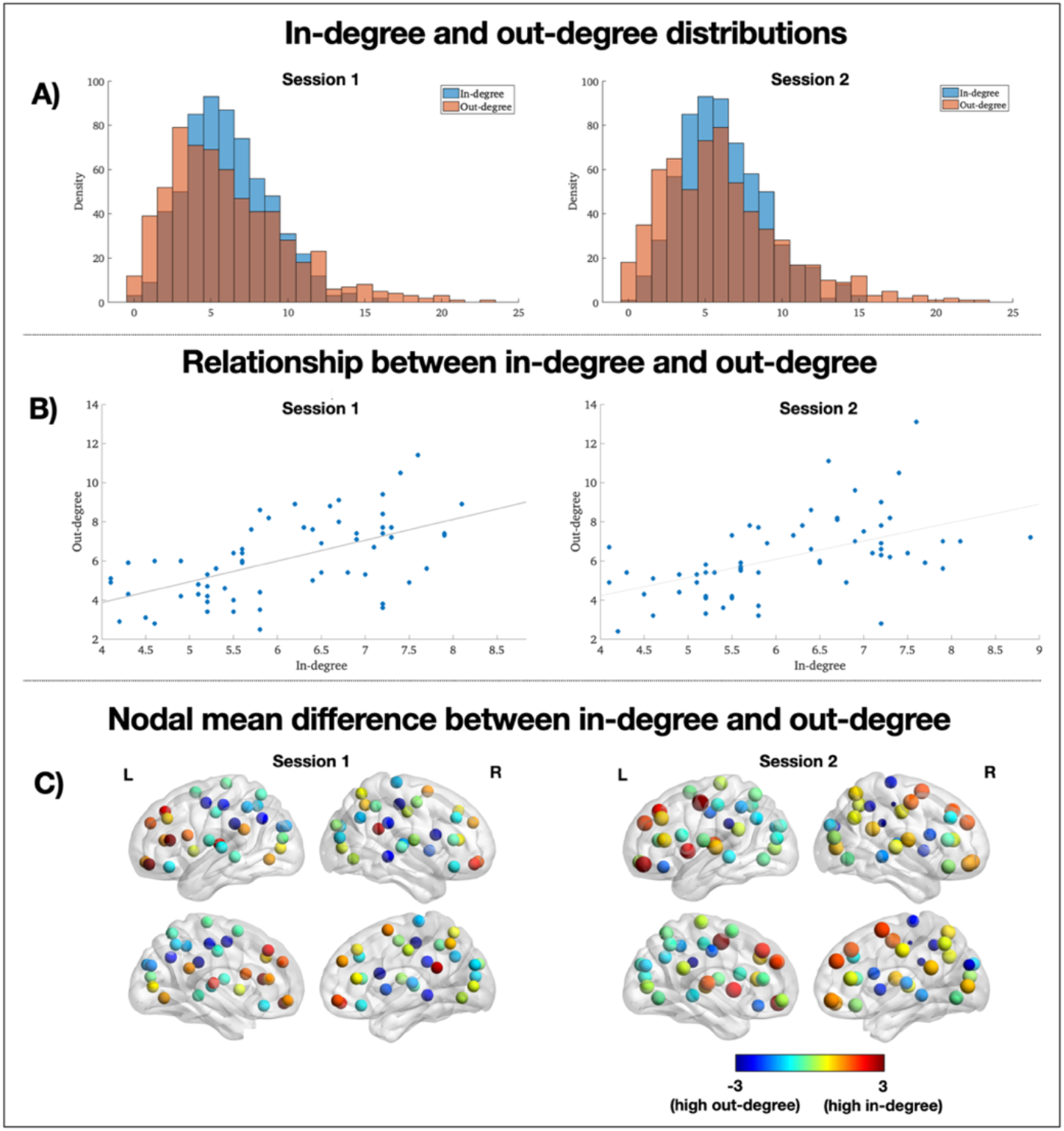
A) Degree distributions of all nodes from all subjects. B) Scatterplot between In- and Out-Degree. Blue dots denote each node (here, 62 nodes), averaged over all subjects. C) Difference maps between In- and Out-Degree for each node, averaged across subjects.

### Network-specific mTE effects for Out-Degree but not In-Degree

Next, we explored In- and Out-Degree for significant mTE connections in six resting-state networks (Yeo et al., 2011), including the visual network (VIS), somatomotor network (SMN), attention network (ATT), frontoparietal network (FPN) and default mode network (DMN). For the limbic system (LIMBIC), only nodes from the orbitofrontal cortex were selected as we did not study deep brain regions in this study. For Out-Degree, we observed a significant difference between networks in session 1: F(5,54) = 8.25, *p* < 0.001. SMN had a greater Out-Degree than VIS (p = 0.004), LIMBIC (p < 0.001) and DMN (p = 0.007), and LIMBIC had a lower Out-Degree than ATT (*p* < 0.001) and FPN (*p* = 0.003). In session 2, a similar trend was seen: F(5,54) = 3.67, *p* = 0.006. SMN had a lower Out-Degree than LIMBIC (*p* = 0.004). Additionally, LIMBIC had a lower Out-Degree than ATT (*p* < 0.001) and FPN (*p* = 0.002). We observed no statistically significant difference between networks for in-degree (F(5,54) = 2.38, *p* < 0.051 – session 1; F(5,54) = 0.95, *p* < 0.454 – session 2). See Figure 3. These findings suggest that specific networks have divergent Out-Degree and, by extension, differences in the rate of network information between networks.

**Figure 3:**
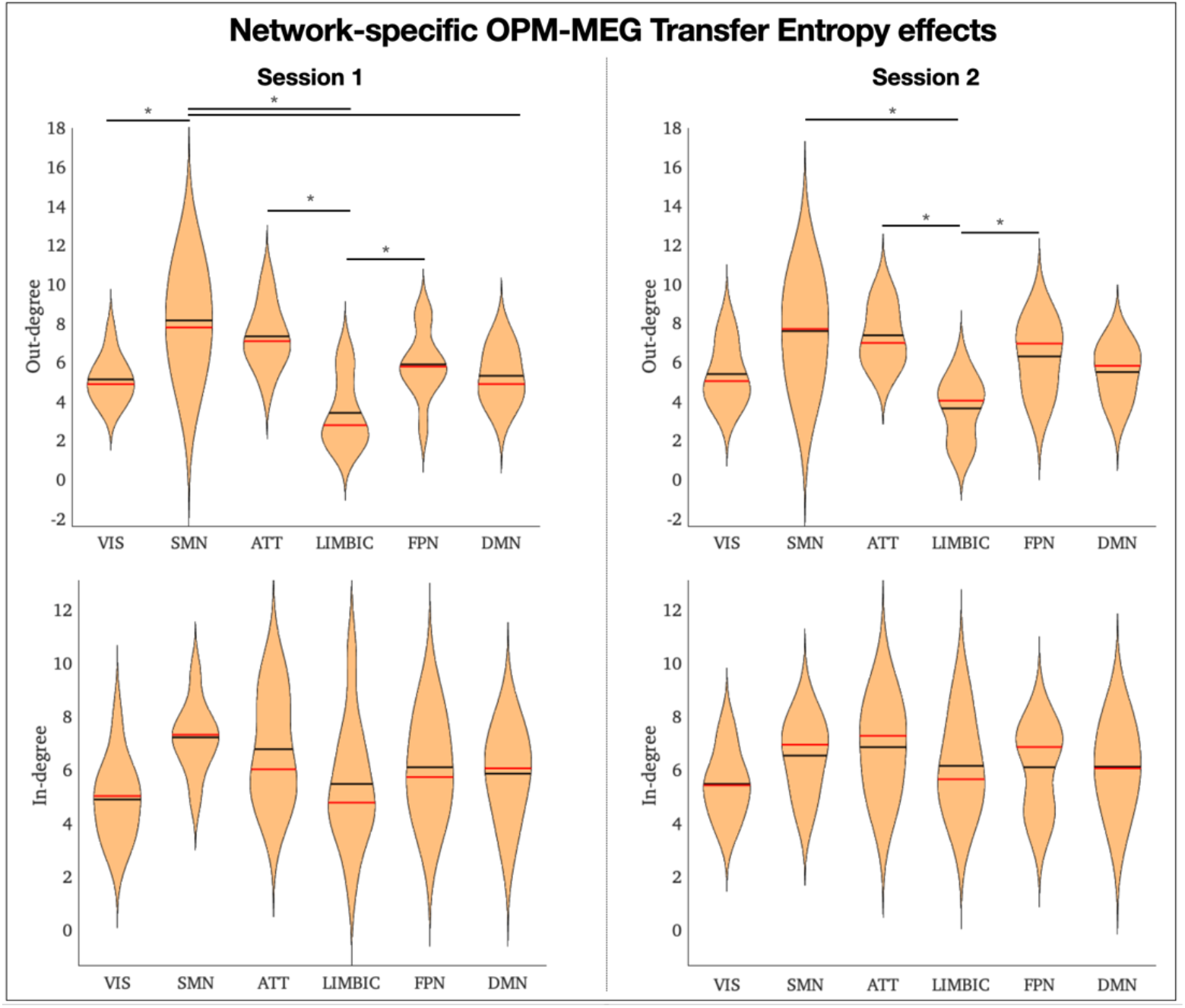
We calculate each subject’s average Out-Degree and In-Degree within each resting-state network. Input values to ANOVA are 6 networks and 10 subjects. Lines indicate statistically significant post-hoc comparisons. Red line = mean; black line = median.

**Figure 4:**
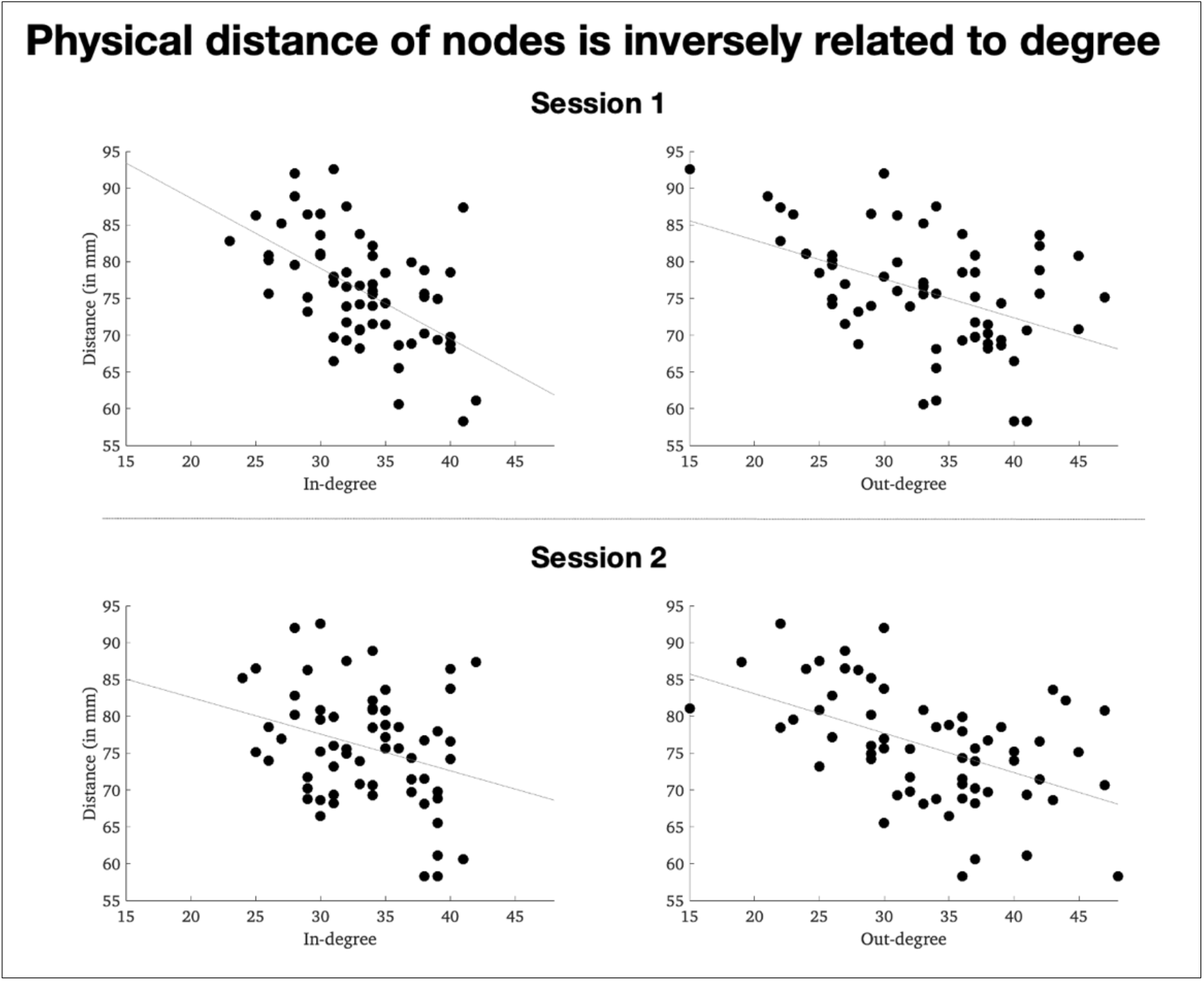
Two session scatter plots between Euclidian distance (y-axis) for each node and In/Out-Degree (x-axis) of in and out-degree, averaged across subjects.

### In- and Out-Degree are inversely related to the physical distance between nodes

After establishing the network-specific effects of mTE, we wanted to examine the relationship between in- and out-degree and physical distance between nodes. To this end, we performed calculations of Euclidean distance between nodes. The results of the node-specific correlation analysis revealed significant findings in both Session #1 and Session #2. We observed a strong negative correlation, Spearman’s ρ(61) = -0.62, p < 0.001 (session 1 – in-degree); Spearman’s ρ(61) = -0.54, p < 0.001 (session 1 – out-degree); Spearman’s ρ(61) = -0.44, p < 0.001 (session 2 – in-degree) Spearman’s ρ(61) = -0.66, p < 0.001 (session 2 – out-degree), indicating that shorter inter-node spatial distances were associated with stronger connectivity.

## Discussion

OPM-MEG is an exciting development in human neuroscience as it provides a wearable design and a high temporal resolution while being less susceptible to volume conduction than EEG (Brookes et al., 2022). This study is a preliminary investigation for directed connectivity studies with OPM-MEG. Firstly, we observed a positive correlation between In- and Out-Degree nodes, indicating that nodes with many incoming connections also have many outgoing connections (Figure 3B). We observed that out-degree, but not in-degree connectivity, displayed a heavy tailed degree distribution where a small subset of nodes have a disproportionally high degree. These highly connected nodes likely serve as brain hubs interconnecting diverging networks, as is common in networks with heavy-tailed degree distributions (Figure 3A – see Heuvel & Sporns, 2011).

Most brain connectivity measures used in human neuroscience are undirected, meaning they do not provide information about the brain directionality (Rubinov & Sporns, 2010). However, TE provides a valuable tool to investigate directed connectivity, enabling us to gain insights into the specific pathways of information transfer between brain regions, in this study, measured by in- and out-degree of nodes. This aspect of TE enhances our understanding of the underlying neural dynamics and mechanisms of communication within the brain. Another strength of mTE is its ability to overcome the issue of zero-lag connectivity, which is often associated with spurious activity and source leakage between local sources in MEG data, leading to misleading connectivity estimations (Brookes et al., 2012). Our findings are similar across two scanning sessions for most brain connections (see Figure 7), consistent with previous work, with a strong mTE correlation between OPM-MEG scans (Rier et al., 2023). We believe more extensive studies are needed to fully confirm connectivity measures’ reliability in OPM-MEG.

Two of our separate findings are worth discussing in combination: i) specific networks, particularly the sensorimotor network, have higher out-degrees than other networks (Figure 2); and ii) high-degree nodes are more spatially proximate than low-degree nodes (Figure 3). Although the relationship between the brain’s structure and function is complex, our findings tie in with previous research showing that sensory cortices are negatively correlated with physical distance between brain regions – i.e., they preferentially have a ‘local connectivity’ pattern (Sepulcre et al., 2010), and also show less variability across subjects (Mueller et al., 2013). These findings suggest that OPM-MEG-directed connectivity can uncover biologically plausible network patterns, but future work combining OPM-MEG and advanced diffusion imaging may further elucidate the brain’s function-structure relationship.

Some limitations need to be considered in our study. A potential concern lies in the necessary band-pass filtering and downsampling of OPM-MEG data in the context of information-theoretic approaches such as mTE or Granger Causality. There is ongoing debate regarding the impact of band-pass filtering on the results obtained from these methods (Barnett & Seth, 2011). Similarly, filtering is contraindicated in TE due to its potential to alter the temporal structure of the data (Weber et al., 2017). In our study, we needed to filter the data between 1 and 48Hz to ensure an acceptable signal-to-noise ratio, considering the narrow bandwidth limitations of the Alkali-based OPM-MEG sensors used in this study. However, we acknowledge that this filtering decision represents a trade-off, and further investigations may be warranted to explore the impact of different filter settings on the TE results. We also deemed that downsampling is needed to ensure that mTE is computationally tractable for OPM-MEG data. However, we carefully considered our temporal downsampling process, ensuring that the order of downsampling was equal to the longest lag used in this study, i.e., 10, to reduce the probability of false negative connections (Weber et al., 2017).

For future studies, we aim to assess the in- and out-degree of other directed connectivity measures allowing for investigation in specific band-pass frequencies. Frequency-specific approaches such as Partial Directed Coherence (Baccalá & Sameshima, 2001) and Directed Phase Lag Index (Stam & van Straaten, 2012) are candidate methods to generate frequency-based directional connectivity matrices.

In conclusion, OPM-MEG research is expected to experience substantial growth, and this research suggests that the use of directed connectivity measures can offer valuable insights into the mechanisms behind brain communication. These measures can enhance our understanding of information transmission and processing within the brain, contributing to advancements in neuroscientific knowledge.

